# *Pseudomonas* virulence factor SaxA detoxifies plant glucosinolate hydrolysis products, rescuing a commensal that suppresses virulence gene expression

**DOI:** 10.1101/2025.04.25.650564

**Authors:** Kerstin Unger, Rebecca Ruiter, Michael Reichelt, Jonathan Gershenzon, Matthew T. Agler

## Abstract

Plants produce a plethora of specialised metabolites that often play important roles in their defence against pathogenic microbes or herbivorous insects. Exposure of leaf colonising microbes to these metabolites influences their growth and we hypothesize that it also has consequences for microbe-microbe interactions and bacterial recruitment to leaves. In Brassicaceae plants like the model plant *Arabidopsis thaliana*, glucosinolates and their biologically active derivatives, the isothiocyanates, are major defence metabolites. Adapted plant pathogens like *Pseudomonas* spp. use the hydrolase SaxA to convert the antimicrobial isothiocyanate sulforaphane to a non-toxic amine, whereas non-adapted commensal microbes are inhibited by this plant toxin. We used *Plantibacter* sp. 2H11-2 as a model commensal in co-culture with either *Pseudomonas viridiflava* 3D9 wildtype or a *saxA*-knock-out mutant. Both strains were isolated from the same wild *A. thaliana* population. Without isothiocyanate, *Plantibacter* grew better alone than with *Pseudomonas*, a potential competitor. At high isothiocyanate concentrations, however, the commensal was dependent on SaxA-mediated isothiocyanate degradation in both solid and liquid medium. At intermediate isothiocyanate concentrations, *Plantibacter*’s transcriptome changed in response to sulforaphane in mono-culture but not in co-culture with *Pseudomonas*, suggesting it was fully protected from this toxin. In return, *Plantibacter* caused transcriptional changes in *Pseudomonas*, suppressing biofilm formation and increasing amino acids metabolism gene expression which might suppress virulence and so contribute to plant health. Together, we find that an antimicrobial plant metabolite can force a commensal to depend on a pathogen-produced virulence factor, but that the interaction overall might benefit the plant by limiting pathogenicity.

## Introduction

Plants are colonized by various microbes, often in a deterministic fashion [1], with the host genotype playing a crucial role in microbial recruitment and community stability [2,3]. The host plant has several systems to control microbial colonizers, including the immune system [4] and production of specialised metabolites [2,5]. Microbial recruitment to the rhizosphere due to root-exuded metabolites has been intensively studied [6,7] but leaves also produce plant metabolites that can shape microbial recruitment [8–10]. For instance, bacterial hub taxa across *Arabidopsis thaliana* accessions were associated with the specialised metabolite 8MSOO-GLS (8-methylsulfinyloctyl-GLS) [2]. Since hub taxa can be centres of interaction networks and influence other microbes, plant specialised metabolites may exert widespread control over microbial interactions [11].

Plant specialised metabolites are probably best known for their antimicrobial effects and defence against pathogenic microbes [12–15]. As a reaction to plant defences some pathogens evolved enzymes to detoxify these metabolites [16–19] which changes the chemical landscape of the leaf. We hypothesize that these could have a large influence on microbial interactions, and thus impact leaf colonisation, but few investigations have covered this topic. Microbial interactions can also be mediated by better known mechanisms, such as competition for nutrients [20], production of antimicrobial substances to inhibit competitors [21], and metabolite cross-feeding [22]. To specifically study the effect of detoxification on leaf microbial interactions, we chose the glucosinolate-myrosinase defence system in *Arabidopsis thaliana* as a model.

Glucosinolates (GLS) are defence metabolites in Brassicacaeae plants like *A. thaliana* as well as various crops, including *Brassica oleracea*, where they are stored in large amounts in leaves [23,24]. GLS themselves have no antimicrobial effect, but they are activated by hydrolytic breakdown catalysed by myrosinases (β-glucosidases) [25]. The most studied GLS breakdown products are isothiocyanates (ITCs), which have antimicrobial activity against diverse pathogenic leaf colonizers [13,17,19,26]. Leaf microbes are exposed to ITCs when pathogens cause necrotic lesions or herbivorous insects bite into the leaf and myrosinases come in contact with GLS [25,27]. Additionally, low levels of ITCs are suggested to be constantly present inside the leaf apoplast because of constant turnover of GLS as part of plant sulphur cycling [28]. Exposure of leaf bacteria to ITCs may also happen on the leaf surface when bacteria metabolize GLSs [8]. Here, we focus on 4MSOB-ITC (4-methylsulfinylbutyl-ITC, sulforaphane) which is the major breakdown product of GLS in the widely used reference *A. thaliana* genotype Col-0. It was shown to reduce bacterial loads of non-adapted *Pseudomonas* spp. or *Pectobacterium* spp. in leaves and inhibits bacterial growth *in vitro* [17,29]. Additionally, 4MSOB-ITC reduces the virulence of plant pathogens like *Pseudomonas syringae* and *Xanthomonas campestris* [30,31].

ITCs represent a major barrier to microbial colonisation, so some well-adapted *A. thaliana* pathogens express *sax* genes (survival in *Arabidopsis* extract) that increase their virulence. These genes encode efflux pumps (*saxF, saxG, saxD*), an uncharacterized protein (*saxB*), a transcriptional regulator (*saxC*) and an ITC hydrolase (*saxA*) [17,32]. The SaxA protein degrades a broad spectrum of ITCs including 4MSOB-ITC to their corresponding amines [32] rendering them non-toxic [17,19]. Although the effects of ITCs on microbes beyond plant pathogens are less studied, they may be based on unspecific modes of action like disturbing protein structures, impairing membrane stability, or general stress responses [33–35]. In a recent study, we demonstrated growth inhibition of several commensal bacterial strains, including *Plantibacter* spp., by 4MSOB-ITC and allyl-ITC *in vitro* [8].

Understanding the interactions between pathogens and commensals is important because commensal bacteria can prevent plant disease by pathogen suppression [36,37], which might help to improve plant health and also increase the effectiveness of biocontrol agents [38,39]. The reverse effect of pathogens on commensals is less well-known but likely as important for understanding the consequences of these interactions. Since pathogens and commensals can be isolated together from healthy *A. thaliana* leaves [1], we hypothesize that plant specialized metabolites like ITCs play a major role in pathogen-commensal interactions with effects on bacterial leaf colonisation and ultimately on plant health. Specifically, SaxA-mediated ITC degradation could serve as a public good that not only benefits the pathogenic degrader strain but also non-degrading commensal bacteria. We investigate this hypothesis (Fig. 1A) and its temporal and spatial implications using the pathogen *Pseudomonas viridiflava* 3D9 (hereafter: Ps), which degrades ITCs via SaxA [8] and was co-isolated from wild *A. thaliana* plants together with the ITC-sensitive, commensal strain *Plantibacter* sp. 2H11-2 (hereafter: Pl) [1].

**Figure 1:**
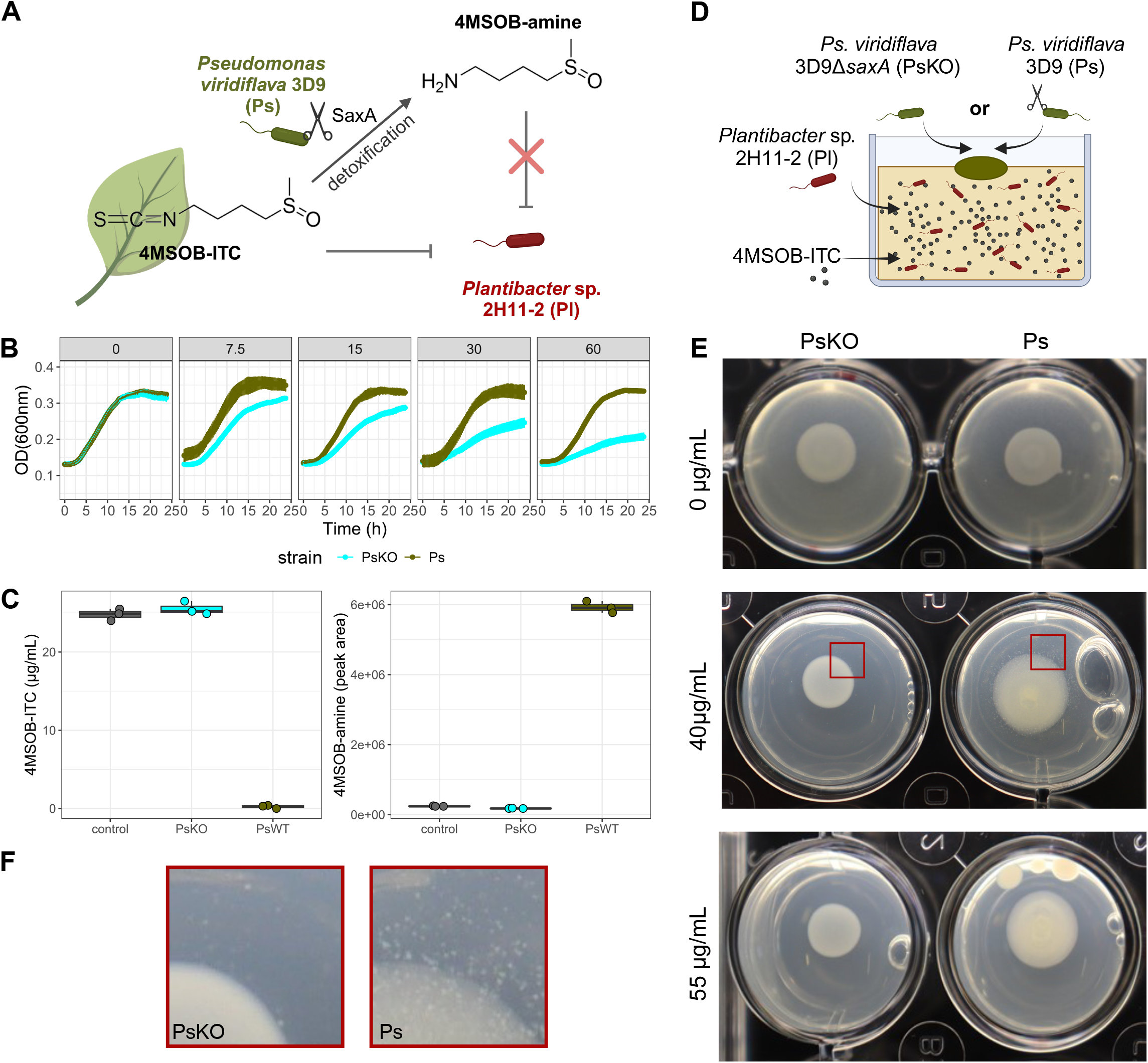
SaxA as public good in the interaction between *Pseudomonas viridiflava* 3D9 (Ps) and *Plantibacter* sp. 2H11-2 (Pl). (**A**) Schematic drawing of the research hypothesis. *Arabidopsis thaliana* Col-0 produces 4MSOB-ITC as defence compound, plant pathogen *Pseudomonas viridiflava* 3D9 (Ps) can degrade it to 4MSOB-amine. The growth of commensal *Planibacter* sp. 2H11-2 (Pl) is inhibited by 4MSOB-ITC but not by 4MSOB-amine. Thus we hypothesize that SaxA-mediated ITC degradation functions as public good which benefits not only Ps but additionally Pl. Created with Biorender.com. (**B**) Growth curves of Ps and PsKO (Ps 3D9Δ*saxA*) with 4MSOB-ITC concentrations from 0 to 60 µg/mL (mean ± standard deviation of n=3 are shown). (**C**) Quantification of 4MSOB-ITC and its breakdown product 4MSOB-amine by LC-MS in supernatants of bacterial overnight cultures (n=3). Non-inoculated R2A broth supplemented with 30 µg/mL 4MSOB-ITC served as control. (**D**) Public good assay: Schematic cross section of the agar plate assay to visualize a public good effect of Ps on Pl in 4MSOB-ITC background. 4MSOB-ITC at different concentrations and Pl cells in low density (OD = 0.001) were mixed in R2A agar and poured in wells of a 24-well plate. 2 µL drop spots of Ps or PsKO were placed on top of the agar layer and growth was observed after four days of incubation at 30°C. (**E**) Growth of Pl (in the agar), and Ps and PsKO (colony in the middle) at different 4MSOB-ITC concentrations. (**F**) Close-up of PsKO/Pl and Ps/Pl (boxes marked in E).

## Material and methods

### Strains and culture conditions

*Pseudomonas viridiflava* 3D9 (Ps) and *Plantibacter* 2H11-2 (Pl) were isolated from wild *A. thaliana* leaves in 2019 [1,8]. In whole genomes, we found homologues of *sax* genes in Ps, but not in Pl. Namely Ps contains one copy each of the ITC-hydrolase gene *saxA*, the transcriptional regulator gene *saxC, saxB* with unknown function, and the efflux systems *saxF* and *saxG*. Additionally we annotated two copies of the efflux system *saxD* [8]. *Pseudomonas viridiflava* 3D9Δ*saxA* (hereafter: PsKO) was generated as part of this study. If not mentioned otherwise, all strains were grown in R2A broth or on R2A agar (yeast extract 0.5 g/L, peptone 0.5g/L, casein hydrolysate 0.5 g/L, glucose 0.5 g/L, soluble starch 0.5 g/L, K_2_HPO_4_ 0.3 g/L, MgSO_4_ 0.024 g/L, sodium pyruvate 0.3 g/L, if applicable: 1.5 g/L agar; pH=7.2±0.2 [40]) at 30°C. Throughout the experiments, we used 4MSOB-ITC (L-Sulforaphane, CAS 142825-10-3, ≥95%, Sigma Aldrich) dissolved in DMSO (Dimethyl sulfoxide, Carl Roth).

### Knock-out of *saxA* in *Pseudomonas viridiflava* 3D9

To knock-out *saxA* in Ps, we used a heterologous recombination protocol [41]. Details are described in the Supplementary Methods.

### Agar-based public good assay

R2A agar was cooled down to 45 °C and Pl overnight culture was normalized to OD=0.01 before mixing it into the warm agar in a final concentration of OD=0.001. 4MSOB-ITC or DMSO was added in concentrations ranging from 0-55 µg/mL, and 1 mL of medium with Pl and ITC was poured into individual wells of a 24-well plate. Ps and PsKO pre-cultures were normalized to OD=0.2 and 2 µL were drop-spotted in the middle of a well as soon as the agar had solidified. The plate was incubated at 30°C for up to 4 days.

### Bacterial co-culture experiments

#### Growth curves and endpoint bacterial loads

Overnight cultures of Ps, Pl and PsKO were normalized to OD_600_ = 0.4 and equal volumes of two strains or one strain and pure R2A broth were mixed to generate the inocula. 90 µL R2A broth with 4MSOB-ITC in concentrations ranging from 0 to 60 µg/mL were pipetted in individual wells of a 96-well plate. 10 µL of normalized bacterial cultures were used as inoculum. Thus, each strain was present at OD_600_ = 0.02 in each well. R2A broth served as non-inoculated control. The plate was sealed with transparent foil to prevent evaporation of the ITC and the OD_600_ was measured every 15 min after 1 min of orbital shaking and recorded using the software i-control 2.0. The raw data was processed with custom scripts in R. After 22 h of incubation, a 10-fold serial dilution was prepared from each sample and 20 µL of up to three dilution steps per sample were spread on R2A plates. For 0-30 µg/mL. it was sufficient to plate the co-cultures on simple R2A plates. For 60 µg/mL, however, the count of Pl was much smaller than the Ps or PsKO count and therefore we plated the co-cultures on selective R2A plates supplemented with 1.5 µg/mL kanamycin where Pl grew but Ps or PsKO were inhibited. Pl counts on R2A+Kan were normalized to counts on R2A to make all conditions comparable.

#### Bacterial loads and 4MSOB-ITC and -amine analysis over time

As described for the growth curves, bacterial overnight cultures were normalized to OD_600_ = 0.4 and mixed in equal amounts or with R2A broth. The cultures were started with 0, 30 or 60 µg/mL 4MSOB-ITC in a total volume of 500 µL. They were incubated at 30°C at 200 rpm and sampled in regular intervals. At each sampling timepoint, 10 µL were taken and a 10-fold serial dilution was prepared in PBS and 20 µL per dilution step and samples were plated on R2A agar. Additionally, 20 µL of the supernatants were mixed with 180 µL MiliQ water and frozen immediately at -20°C for 4MSOB-ITC and -amine quantification.

### Quantification of 4MSOB-ITC and breakdown products with LC-MS

4MSOB-ITC and 4MSOB-amine were measured in the supernatants of bacterial cultures. To confirm the successful knock-out of *saxA* in PsKO and to follow the degradation of the ITC over time. All supernatants were stored at -20°C before the measurement. Freshly prepared external standard curves of 4MSOB-ITC (L-Sulforaphane, CAS 142825-10-3, Sigma Aldrich) and 4MSOB-amine (4-methanesulfinylbutan-1-amine, Enamine Germany GmbH, Frankfurt a.M., Germany) were measured together with the samples. The measurements were conducted as described in [8] and details are described in the Supplementary Methods.

### Transcriptomics

#### Growth of bacterial cultures

Overnight cultures were normalized to OD600=0.2 and mixed in equal amounts with either a partner strain (co-cultures) or R2A (mono-cultures). 100 µL were used as inoculum for 1.9 mL R2A supplemented with 15 µg/mL 4MSOB-ITC or DMSO as a control. The cultures were incubated in 5 mL tubes on a shaker at 30°C, 150 rpm, for 17-18 h. Mono-cultures and co-cultures were performed in separate experiments.

#### RNA extraction

Bacterial cultures were harvested by centrifugation at 5000 x g for 20 min at 4°C. Supernatants were discarded and the cell pellets were frozen in liquid nitrogen. Total RNA was extracted using lysozyme and a hot phenol extraction protocol followed by a DNAse treatment, details are described in the Supplementary Methods. All samples were sent to Eurofins Genomics (Konstanz, Germany) on dry ice, where the rRNA depletion, library preparation and sequencing were performed.

#### Data analysis

Raw reads were trimmed using the implementation of trimmomatic in the GREP2 package in R [42] and were then mapped to the whole genomes of Pl and Ps [8] using Rsubread function align [43] and assigned to genes using the featureCounts function [44]. Low abundant genes were excluded from the subsequent analysis and only genes with 10 reads in at least 3 samples were considered. The count data of the remaining genes was normalized and Log2FoldChanges (L2FC) were calculated using DESeq() function with default settings in the DESeq2 package which include a Wald test and Benjamin-Hochberg correction [45]. Using the ggplot2 package [46], a PCA was plotted of the top 500 genes with the highest variance using rlog-transformed data for the plotPCA function in DESeq2 package. To analyse differentially expressed genes (DEGs) the results of pairwise DESeq2 calculations were shrunk using lfcshrink function with apeglm [47]. After checking the dispersal in MA plots, the data was subset to keep only significant (p < 0.05) genes with |L2FC| > 1 for the analysis. Venn diagrams with shared and unique DEGs between treatments were determined and visualised using the package VennDetail [48]. We summarized Pl’s reaction to the ITC by filtering for DEGs with |L2FC|>2 to prioritize important processes. To characterize the reaction of Ps or Pl to the partner strain, we generated gene to GO term maps for Pl and Ps with GO annotations of their whole genomes which were annotated using bakta [49]. Next we tested for significant enrichment of GO terms using the function runTest with the classic algorithm followed by a Fisher test using the packages topGO [50]. For each pairwise comparison significantly enriched GO terms were determined. The package VennDetail [48] was used to assess shared and unique terms between treatments. GO terms of each strain’s reaction to the partner with or without 4MSOB-ITC were plotted together.

## Results

### SaxA-mediated ITC degradation by the pathogen *Pseudomonas viridiflava* (Ps) facilitates growth of the co-colonising commensal *Plantibacter* (Pl)

To investigate a potential positive effect of SaxA-mediated 4MSOB-ITC degradation on ITC-sensitive commensals (Fig. 1A), we generated a knock-out of *saxA* (hereafter: PsKO) in our model pathogen strain *Pseudomonas viridiflava* 3D9 (hereafter: Ps). We established earlier that the wildtype strain degrades 4MSOB-ITC to 4MSOB-amine [8]. The knock-out of *saxA* resulted in a complete loss-of-function, showing that SaxA in Ps is completely responsible for the degradation of 4MSOB-ITC (Fig. 1C). Although homologs of ITC efflux pumps were identified in the genome of Ps [8], growth of PsKO was significantly inhibited by increasing 4MSOB-ITC concentrations at 30 °C (Fig. 1B). To check the interaction between Ps or PsKO with Pl we established a “public good assay” (Fig. 1D): Pl was poured into R2A agar together with different concentrations of 4MSOB-ITC and Ps or PsKO were drop spotted on top of the agar. At 0 µg/mL 4MSOB-ITC, Pl grew densely in the entire well (opaque agar) with both Ps strains. At a high ITC concentration of 55 µg/mL, Pl growth was completely inhibited after four days (clear agar). At intermediate 4MSOB-ITC levels (40 µg/mL), however, we observed a zone of Pl colonies around the Ps but not the PsKO colony (Fig. 1E,1F). Taken together, Pl benefits from SaxA-mediated 4MSOB-ITC degradation and is able to grow in the vicinity of Ps.

### The public good effect of SaxA increases in a concentration- and time-dependent manner

In the next step we studied the growth dynamics of the individual Pl, Ps and PsKO strains and their co-cultures in liquid medium supplemented with 4MSOB-ITC concentrations from 0 to 60 µg/mL (Fig. 2A). As observed earlier [8] Pl in mono-culture was clearly reduced by 30 µg/mL and its growth slowed further to only about 1/3 of the original OD at 60 µg/mL 4MSOB-ITC (Fig. 2A). Because growth curves of co-cultures do not allow the differentiation of individual strains, we determined the bacterial counts of Ps, PsKO and Pl after 22 h by plating (Fig. 2B). As expected, Pl counts in mono-culture were reduced with increasing ITC concentration. When no or only little (7.5 µg/mL) 4MSOB-ITC was added, Pl grew better alone than with Ps or PsKO, likely due to higher nutrient availability in the absence of a competitor (Fig. 2B). Since Pl was not completely outcompeted in the presence of Ps there is probably no direct inhibitory action from Ps towards the commensal. At low to intermediate concentrations (15 and 30 µg/mL) there were no significant differences between final bacterial counts in Pl mono-culture and co-cultures. However, at the highest concentration (60 µg/mL) we clearly observed a public good effect and Ps with SaxA but not PsKO rescued Pl growth (p=0.0003 Ps+Pl vs. Pl, p=0.003 Ps+Pl vs. PsKO+Pl; Fig. 2B). Pl growth with PsKO was similar to Pl mono-culture suggesting that the ITC effect dominated, and nutrient competition played only a minor role, if any, at this concentration (Fig. 2B). Because ITC degradation by Ps was already evident after 3-6 h (Supplementary. Feig. 1A,1B), we investigated the interaction between Ps and Pl at these early timepoints. We expected Pl to benefit especially right after complete ITC degradation when it did not pay the cost of SaxA production in contrast to Ps. We chose 30 µg/mL 4MSOB-ITC because Pl was already inhibited at this concentration early in the time course when grown alone (Fig. 2A). Indeed, we observed a public good effect in the Ps+Pl co-culture after 6 h (Fig. 2D, p=0.023 for Ps+Pl vs. PsKO+Pl). However, this effect was not observed after 7.5 h (Fig. 2D) or after 22 h (Fig. 2B) indicating only a transient effect on Pl growth at intermediate ITC levels. Interestingly, at 30 µg/mL ITC Ps also benefited from Pl for a short time between 12.5 and 15 h independent of the ITC content (Fig. 2C) indicating possible mutualistic benefits rather than Pl simply exploiting Ps degradation of the toxin. Taken together, Pl growth was measurably inhibited by intermediate ITC levels (30 µg/mL) but with Ps it can overcome this inhibition. In co-culture at high ITC levels (60 µg/mL) Pl was dependent on SaxA-mediated degradation of 4MSOB-ITC by Ps.

**Figure 2:**
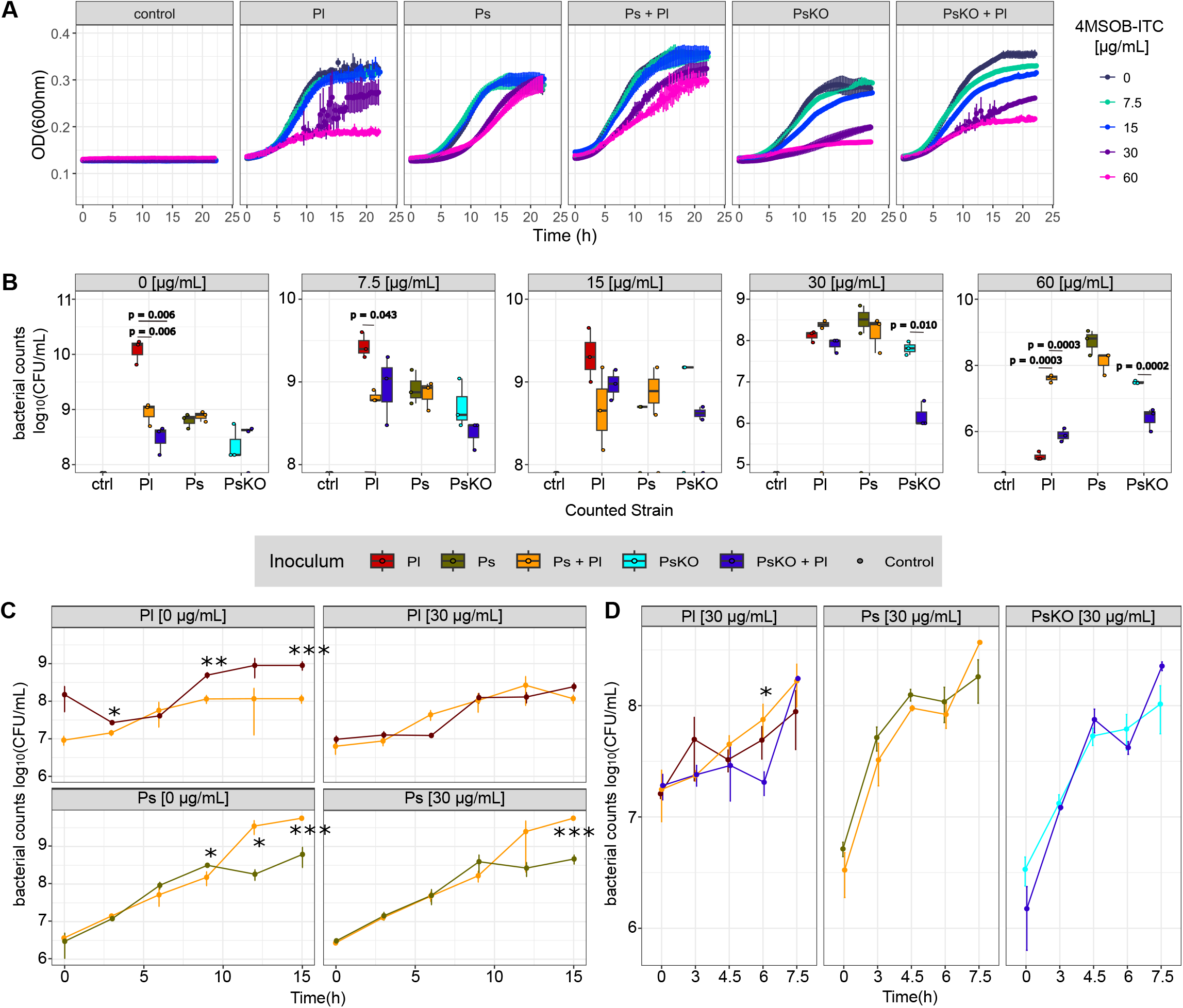
Benefit of Pl from *saxA*-mediated 4MSOB-ITC degradation by Ps over time. (A) Growth curves of mono- and co-cultures of Ps, Pl, and PsKO at 4MSOB-ITC concentrations from 0-60 µg/mL at 30°C (mean ± standard deviation, n=3). The strains were inoculated with OD=0.2, the initial inoculum of the co-cultures was OD=0.4. The OD_600_ was measured in 15 min intervals over the course of 24 h. Control wells were inoculated with sterile medium. (**B**) Bacterial counts of mono- and co-cultures of Ps, PsKO and Pl after 22 h of incubation at 30°C (n=3). P-values only for significant comparisons are shown (p<0.05). (**C**) Bacterial counts of Ps and Pl in mono- and co-cultures with 0 or 30 µg/mL 4MSOB-ITC over the course of 15 h (mean ± standard deviation, n=3). Facets indicate the counted strain and the starting concentration of 4MSOB-ITC. Colours visualize the inoculum as in B. Significant comparisons at all timepoints are shown as asterisks * p<0.05, ** p<0.01, *** p<0.001. (**D**) Bacterial counts of Ps, PsKO and Pl in mono- and co-cultures over the course of 7.5 h (mean ± standard deviation, n=3). Colours, facets and asterisks as shown in (B).

### Low 4MSOB-ITC levels only affect the transcriptome of Pl growing without Ps

Low ITC concentrations are likely to be encountered by bacteria even in healthy leaf tissue [28] where they may not exert significant cytotoxic or growth inhibiting effects [30,31,51] but act on a transcriptional level. In accordance, we did not observe significant growth reduction of Pl when supplemented with 15 µg/mL 4MSOB-ITC (Fig. 2A). To investigate the effects of low 4MSOB-ITC levels on the interaction between Ps and Pl, we grew them either alone or together supplemented with 15 µg/mL 4MSOB-ITC and evaluated their gene expression profiles using RNA sequencing after 17-18 h.

Because Ps protects itself by both detoxification and metabolite export, as expected, its transcriptome was hardly affected by 4MSOB-ITC (Fig. 3A). We detected only 3 significantly differentially expressed genes (DEGs) in Ps (log_2_fold-change |L2FC|>1, p<0.05) in response to the ITC in mono- or co-culture (Suppl. File with all DEGs in all comparisons). At the sampling timepoint, Ps had already been growing without the ITC for 12-15 h due to the rapid rate of degradation. Even though the ITC was no longer present, both *saxA* and *saxB* genes were still slightly but significantly upregulated in response to 4MSOB-ITC in both Ps mono- and co-cultures (Fig. 3B, Suppl. Table 3; *saxA*: L2FC=0.5238, padj=0.0012 (mono); L2FC=0.6296, padj=0.0057 (co), *saxB*: L2FC=0.3690, padj=0.0217 (mono); L2FC=0.4415, padj=0.0451 (co)).

**Figure 3:**
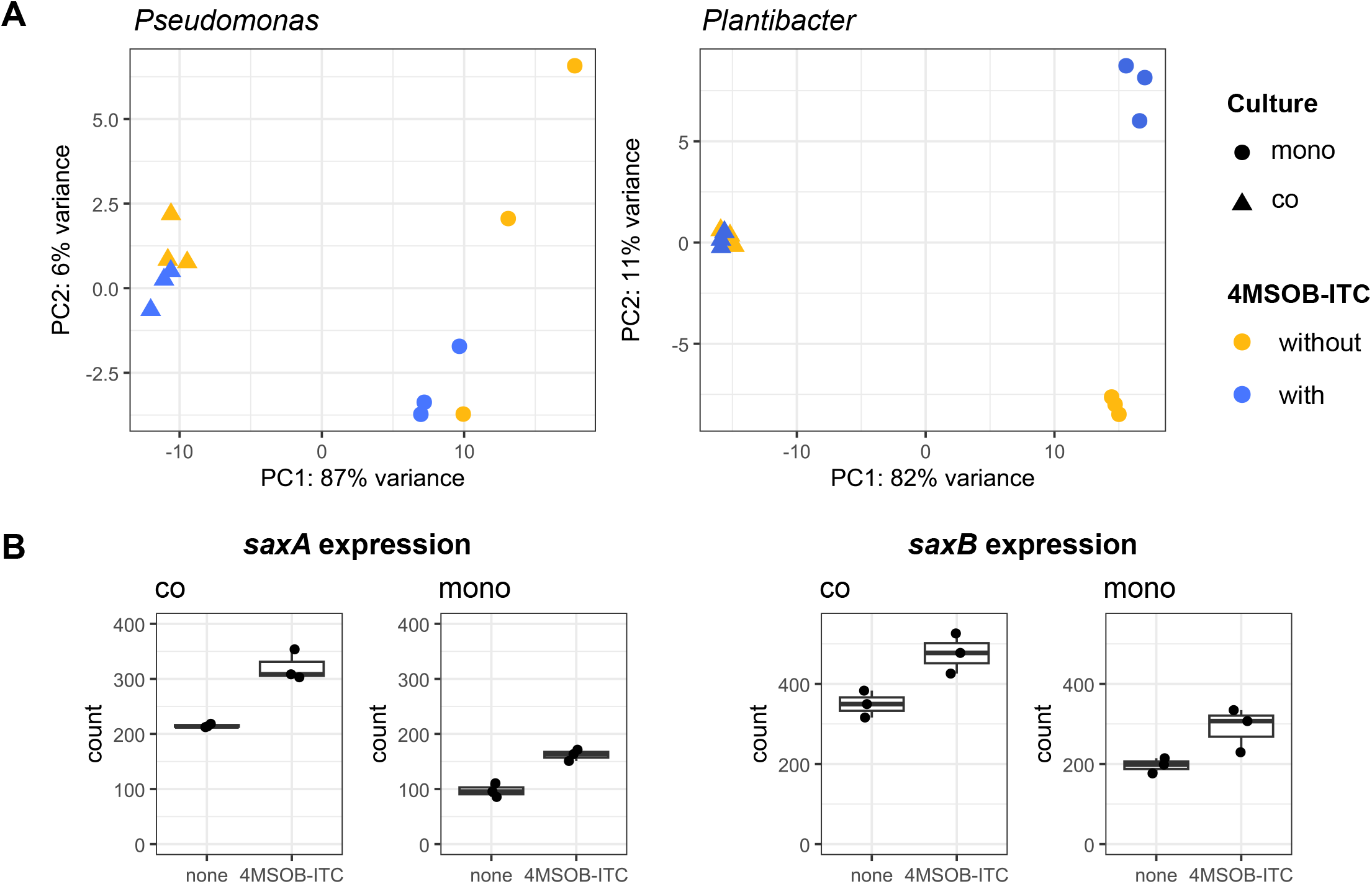
ITC effects on Ps and Pl gene expression, especially *sax* expression. (A) Comparison of partner and ITC effect on both isolates. PCA of rlog-transformed DESeq results. The partner effect on Ps and Pl reads is correlated to a higher proportion of the variance than the ITC effect. Triangles show co-cultures, circles show mono-cultures of Ps (left panel) and Pl (right panel) with or without 4MSOB-ITC (n=3 per condition). (B) Significant upregulation of *saxA* and *saxB* in Ps mono- and co-culture in response to 4MSOB-ITC exposure.

The transcriptional regulator *saxC* was not upregulated upon ITC exposure (Suppl. Table 3) and efflux pumps of type “Multidrug efflux pump subunit AcrB” with COG0841 number which were earlier associated with *saxF* [17,52] were not significantly differentially expressed either (Suppl. Table 4), indicating ITC degradation rather than ITC efflux to be the major resistance mechanism in this strain.

In contrast to Ps, Pl was more affected by the low levels of 4MSOB-ITC. In mono-culture, we detected 100 DEGs (|L2FC|>1, padj<0.05, Suppl. File with all DEGs, Suppl. Fig. 2) and transcriptomes of Pl mono-cultures with or without ITC clearly separated in the PCA (Fig. 3A). A more stringent filtering (|L2FC>2|) left only 17 DEGs (Suppl. Tab. 5). Of these, five were downregulated, and their predicted protein annotations suggest links to costly metabolic processes. Among the 12 upregulated DEGs, two were annotated with proteins which can be involved in oxidative stress responses in other bacteria (ACEBMG_0398, nitroreductase; ACEBMG_15290, NADPH:quinone reductase) [53,54] and another three were annotated as transcriptional regulators (ACEBMG_16910, HTH-type transcriptional regulator CmtR; ACEBMG_17225, HxlR family transcriptional regulator; ACEBMG_03025, AraC family transcriptional regulator) (Suppl. Tab. 5). Taken together, Pl displays a general transcriptional reprogramming after exposure to 4MSOB-ITC with increased defence against oxidative stress and reduced metabolism possibly to save energy. None of these effects were observed when Pl was co-cultured with Ps (Fig. 3A) where no DEGs were identified (padj<0.05, |L2FC|>1), indicating that the Ps SaxA activity fully protected Pl from ITC toxicity at low 4MSOB-ITC levels.

### Co-cultivation with Pl reduces virulence expression in Ps independent of 4MSOB-ITC

Next, we evaluated whether the effect of the Pl or Ps strain on each other went beyond ITC (Fig. 3, Fig. 4A,4B). Interestingly, the reaction of each strain to the other was at least partly influenced by previous ITC exposure even though we only saw a strong ITC effect on Pl mono-cultures where it was not degraded by Ps. There were 565 and 519 DEGs identified in Pl in response to Ps in the presence and absence of 4MSOB-ITC, respectively. More than half of these DEGs were independent of previous ITC exposure (298 DEGs, Fig. 4A). For Ps it was similar, with 210 or 340 DEGs found in response to Pl when the ITC was added or not, respectively. A high fraction (181 DEGs) was independent of 4MSOB-ITC, but without ITC Ps differentially regulated an additional 159 genes in response to Pl. In the presence of ITC, Ps differentially expressed only 29 genes (Fig. 4B), suggesting that Ps responded more to Pl when there was no need to degrade 4MSOB-ITC.

**Figure 4:**
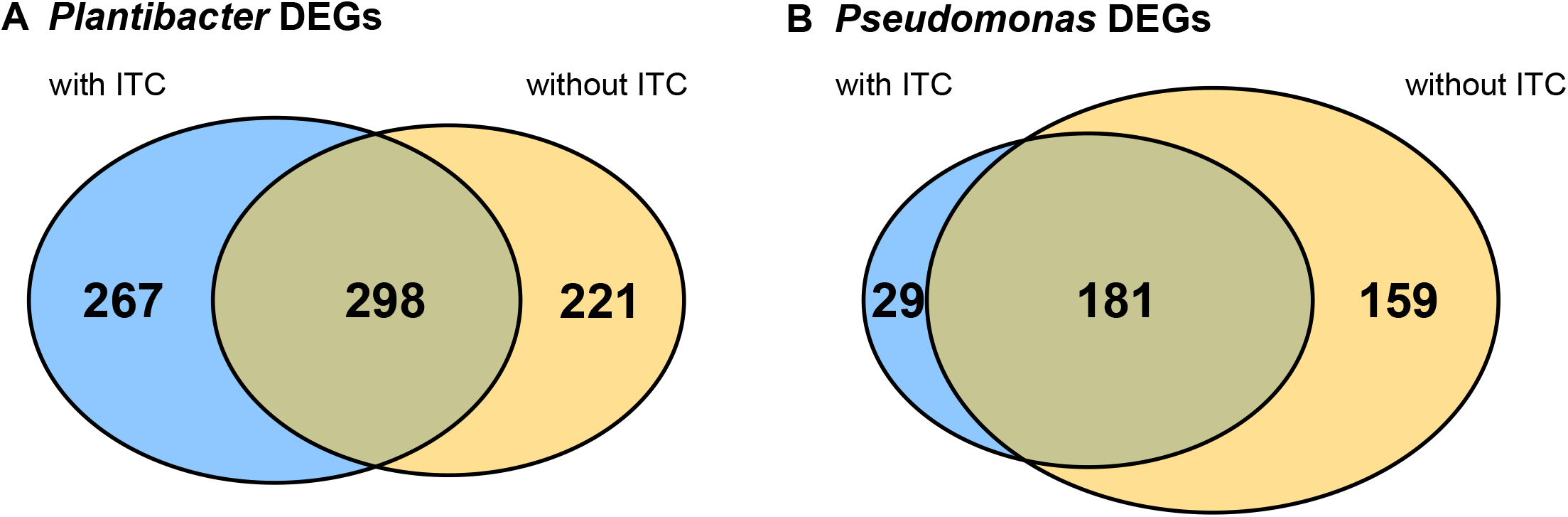
Effect on Pl and Ps on gene expression in each other in relation to previous 4MSOB-ITC exposure. (**A)** Pl reacted to Ps and significantly changed the regulation of 267 genes (L2FC > 1, p < 0.05) when the co-culture was exposed to 4MSOB-ITC, or 211 genes without ITC. 298 DEGs represent Pl’s reaction to Ps independently of 4MSOB-ITC. (**B**) Ps reacts to Pl with 181 DEGs which are independent of previous ITC exposure. This strain regulates more unique DEGs (159) without exposure to 4MSOB-ITC than after exposure (29 DEGs).

In line with this, most processes that were significantly enriched in our GO term analysis were independent of ITC exposure. The most significant (p<0.01, Fisher test) GO terms for upregulated DEGs by Ps in response to Pl (Fig. 5A) were linked to flagellum-dependent motility, chemotaxis and locomotion, catabolism of diverse amino acids (especially Phe, Tyr, Val, aromatic and branched amino acids) and iron transport, all independent of the ITC. Without 4MSOB-ITC Ps in addition upregulated transport of ions like molybdate, while with 4MSOB-ITC, Ps upregulated oxidative stress responses. The downregulated GO terms (Fig. 5B) for Ps without ITC were associated with biosynthesis of alginate, polysaccharides and carbohydrates in general, transmembrane transport processes, protein secretion by the type II secretion system, and processes linked to general protein localization and protein secretion, all of them independent of the ITC. With 4MSOB-ITC, Ps downregulated energy/electron coupled transmembrane transport and ion transmembrane transport processes. The ITC-independent changes suggest a shift in Ps lifestyle from cell aggregates/biofilms to more motile growth in reaction to Pl.

**Figure 5:**
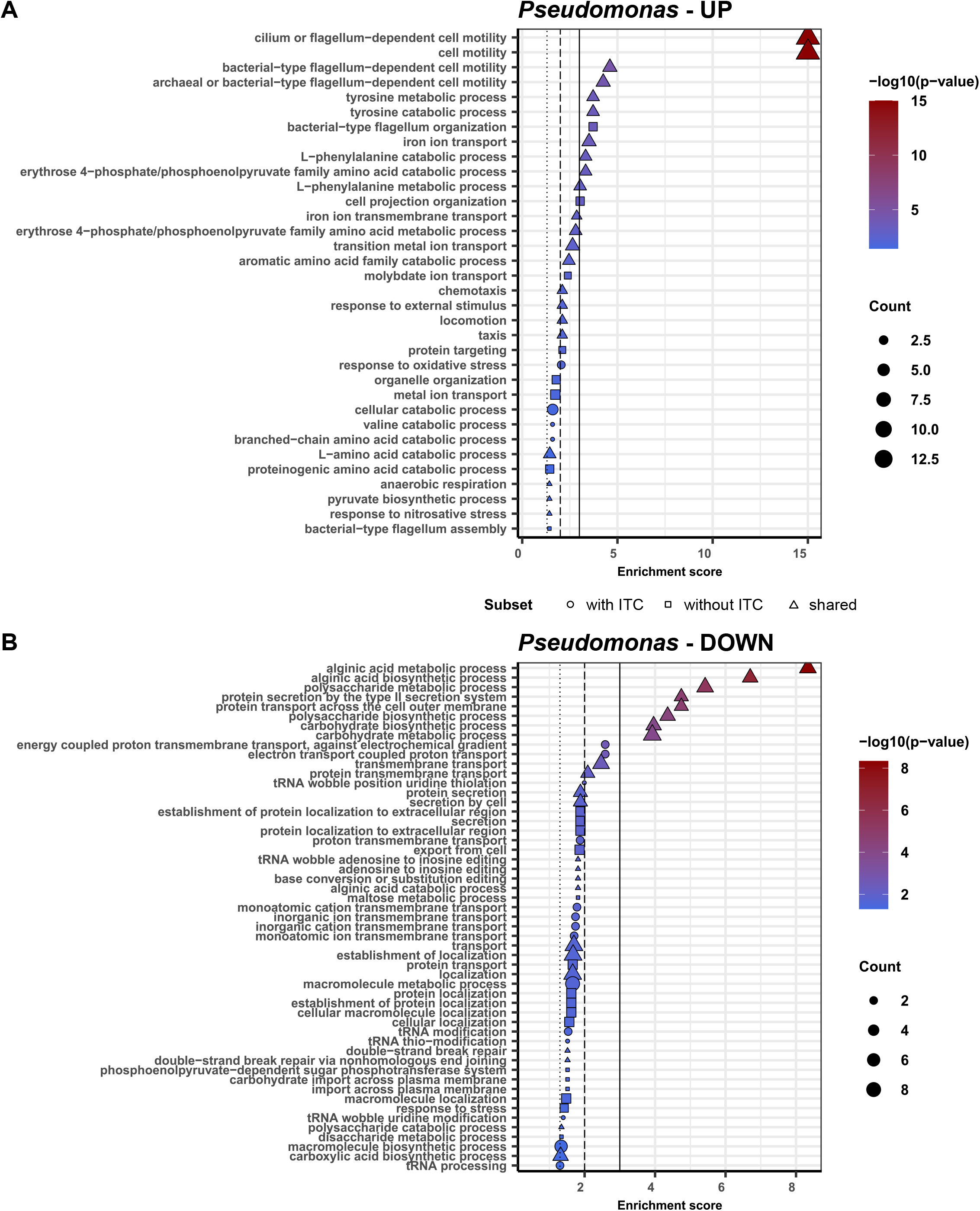
GO term enrichment analysis for the reaction of Ps to Pl with or without ITC. Significantly enriched GO terms for upregulated **(A)** and downregulated (**B**) Ps genes in reaction to Pl. Shared and unique genes are marked by shape, the colour indicates the p-values (Fisher’s significance test), the size correlates to the count number of GO terms for each category. Horizontal lines indicate significance levels: p=0.001 (straight line), p=0.01 (dashed line), p=0.05 (dotted line).

At the same time, the GO processes upregulated by Pl in reaction to Ps all were independent of 4MSOB-ITC. These included catabolic processes related to amino acids, carboxylic acids and organic acids, as well as translation (Suppl. Fig. 3). Pl reacted to Ps by downregulating processes associated with GO terms on transport (metal ions, especially cobalt, magnesium, potassium) which was independent of 4MSOB-ITC. With ITC, Pl additionally decreased carbohydrate transport processes and without ITC Pl additionally decreased processes relevant for ion homeostasis (Suppl. Fig. 3).

In conclusion, the reaction of both strains to each other was mostly independent of the presence of 4MSOB-ITC, likely because of rapid ITC degradation by Ps in co-cultures but suggests that the interaction of Ps with a commensal can shape its virulence.

## Discussion

In this study we showed that plant metabolites can modulate interactions between commensal and pathogenic bacteria. Specifically, detoxification of antimicrobial plant metabolites by one leaf bacterium will directly benefit other sensitive bacteria in its neighbourhood. This public good effect of toxin degradation has so far only been studied in antibiotic degraders [55–57]. Enzymes that degrade antibiotics or catalyse reactions that make them non-toxic can confer a benefit on susceptible co-colonising bacterial strains [58]. This effect is also called “passive resistance” and has been especially studied in connection with β-lactamases that degrade antibiotics like ampicillin or penicillin [55,58,59]. Here we extend this principle to the metabolism of plant-derived antimicrobials.

In the specific case of ITCs, studied here, there are at least two scenarios in which ITC detoxification by SaxA might become relevant for a commensal like *Plantibacter* sp. (Fig. 6). First, when a plant is under attack by an herbivore or pathogen, it may suddenly release high ITC concentrations [27,60]. Second, ITCs may be present in low concentrations in leaf tissue due to constant degradation of GLS to ITCs and cysteine [28] or when GLS-utilising bacteria form ITCs from GLS present in the plant or on the plant surface [8]. Unfortunately, studies measuring ITC concentrations in living plant leaves are rare. Wang et al. [30] measured about 42 µM (= 7.4 µg/mL) 4MSOB-ITC in apoplastic fluid of non-infected *A. thaliana* Col-0 leaves, and we and others [8,61] measured glucosinolates on the leaf surface which can be broken down by some leaf colonisers [8] and likely expose co-colonising strains to ITCs. However, growth in biofilms or cell aggregates [62,63], patchily distribution of microbes [62] and non-uniform distribution of metabolites in leaves [61,64] make it hard to estimate the exposure of individual cells to 4MSOB-ITC. Individual microbes might be locally exposed to even higher concentrations than 42 µM. Our findings illustrate the need to more precisely quantify bacterial exposure to ITCs and other specialised metabolites in living plant tissues.

**Figure 6:**
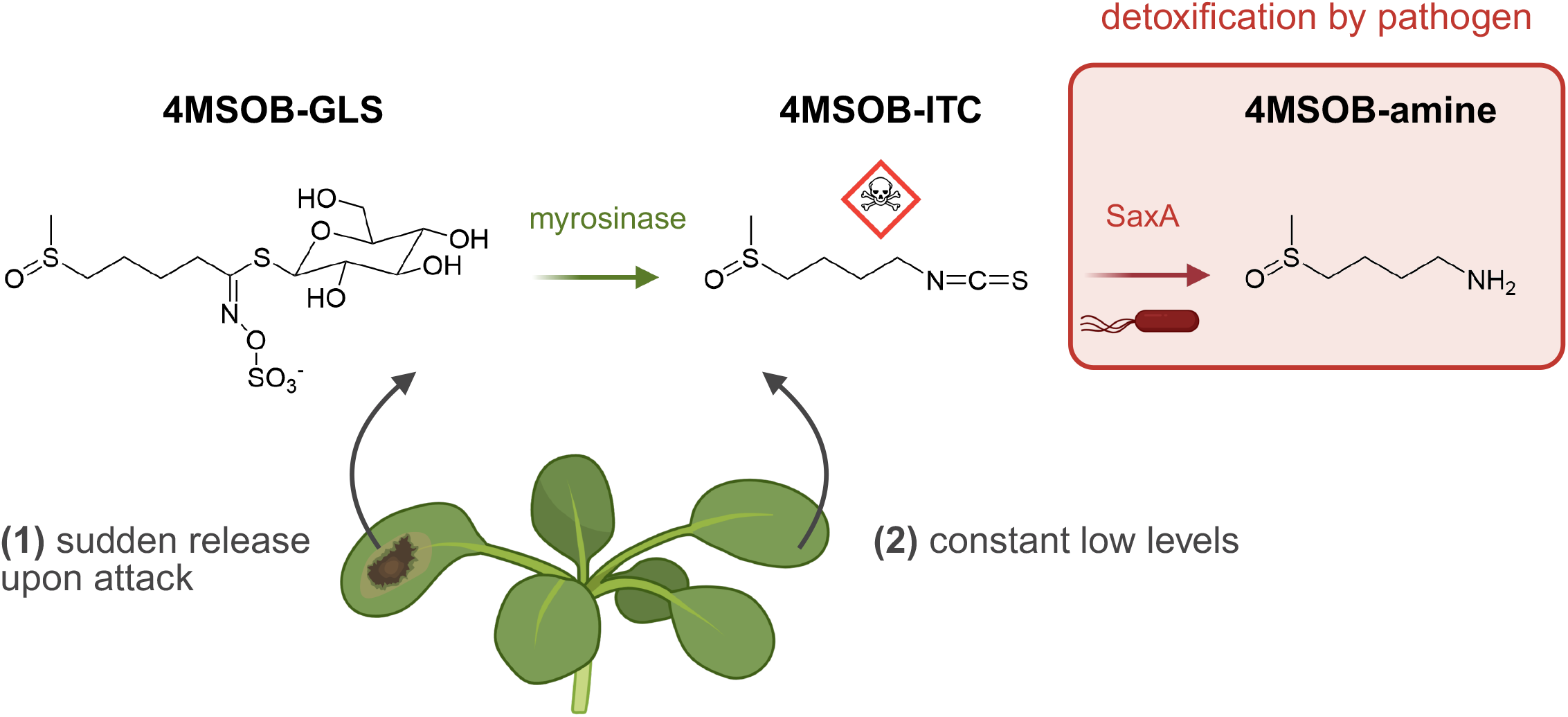
Microbial leaf colonisers are exposed to ITCs. Either when (1) glucosinolates (GLS) are converted to antimicrobial isothiocyanates (ITCs) by myrosinases either in necrotic leaf tissue upon attack by a plant pathogen or when insects bite into a leaf, or (2) when GLS are constantly turned over to ITCs (and cysteine) in low amounts. Plant pathogens like *Pseudomonas viridiflava* 3D9 detoxify ITCs by the ITC hydrolase SaxA to the corresponding amine. The figure shows 4-methylsulfinylbutyl glucosinolate (4MSOB-GLS) which is the major GLS in *A. thaliana* Col-0 and its breakdown products. The figure was created with Biorender.com.

Especially in the first scenario (Fig. 6), the sudden release of high ITC levels presents a serious threat for the fitness of ITC-sensitive commensals, because they severely impair cell membranes and non-specifically bind to proteins, affecting their activity [33,34]. Reductions in the populations of specific commensal bacteria might alter the balance of the entire leaf microbiome, which is of great importance because they are needed to help protect the plant from disease [3,36]. Effects of ITCs on microbial communities were demonstrated in other environments, like in soil amendments where the addition of ITCs shifted the composition of bacterial and fungal communities [65,66]. It seems plausible that sudden high ITC concentrations can also change the composition of the microbial community in leaves because leaf commensals are also inhibited by ITCs [8]. An altered community composition may influence for example its members’ production of secondary metabolites [67] and hence its interaction with the plant host. Therefore, the detoxification of ITCs by SaxA might provide a stabilising effect on the leaf microbiome during pathogen or herbivore attacks, which could benefit the plant. In other words, the well-known role of SaxA as a contributor to pathogen virulence needs to be understood considering its ecological role in a natural leaf bacteriome setting.

In the second scenario, constant low ITC levels in leaves might not directly inhibit bacterial growth but still could alter transcriptional activity of commensals. In our study Pl growth was not inhibited by 15 µg/mL 4MSOB-ITC, but its transcriptome already changed at these low levels. We found several transcriptional regulators to be more expressed when exposed to the ITC, suggesting a general transcriptional reprogramming. Additionally, Pl upregulated genes which were associated with functions in oxidative stress response in other bacteria [53,54]. This agrees with other studies showing effects of sublethal ITC concentrations on bacterial activity like induction of stringent response [68], inhibition of quorum sensing [51] or suppression of virulence [30,31]. In co-culture Ps degraded the ITC present and so Pl’s transcriptome was not affected, demonstrating a clear public good effect even at low ITC concentrations. Nevertheless, previous ITC exposure influenced how Ps and Pl reacted to one another even when the ITC had already been degraded hours before. Thus, changes in the transcriptome may cause subsequent changes in microbial interactions.

Bacterial strains that do not pay the cost of a public good often gain an advantage and may eventually outcompete the producer of the public good, known as the ‘tragedy of the commons’ [69]. We expected to observe that Pl would come to dominate the co-culture because it benefited from SaxA-mediated ITC degradation without having to invest energy. However, we only observed this effect transiently at 6 h at intermediate ITC concentrations (30 µg/mL). In the long run, neither Ps nor Pl outcompeted the other strain. Instead, our experiments demonstrate a clear stepwise benefit for Pl from Ps with increasing ITC concentration until Pl was fully dependent on Ps at 60 µg/mL 4MSOB-ITC. In return, we also observed a transient benefit for Ps from Pl at 30 µg/mL 4MSOB-ITC between 12.5 and 15 h but not after 22 h. This might indicate mutualistic benefits between both strains that can stabilize an interaction [70,71] and therefore may provide an explanation why Ps did not outcompete Pl. At low ITC concentrations (15 µg/mL), the situation is different: each strain upregulated amino acid catabolism in reaction to the other, likely a sign of competition. This fits previous studies that suggested substrate competition to be an important driver of leaf bacterial interactions [20,72]. However, the upregulation of amino acid catabolism in both strains might also be an indication of cross-feeding, but we cannot exclude this possibility without further experiments. Cross-feeding could also be supported by the release of 4MSOB-amine which would likely only be relevant under nitrogen-deficient conditions. Together, we found suggestions for both cooperation and competition between Ps and Pl and the outcome of their interaction was highly depended on the 4MSOB-ITC concentration.

In the absence of ITCs, we observed a clear change in the lifestyle of Ps when it was co-cultured with Pl. Ps increased its motility, decreased biofilm-related transcriptional processes like alginate and polysaccharide production, and protein secretion by the type II secretion system. Alginate production is a major virulence factor of plant pathogenic pseudomonads [73] and the type II secretion system plays a role for virulence of opportunistic *Xanthomonas* sp. [74]. Thus, co-colonisation with a commensal like Pl could confer disease protection to the host plant by suppressing these processes. Interestingly, the reaction of Ps to Pl has similarities to the transcriptome of pathogenic *Pseudomonas syringae* during its epiphytic non-virulent life stage which is characterized by flagellar motility, chemotaxis and higher activity of amino acid catabolism than in its endophytic stage [75]. Moreover, since virulence of Ps relies on nutrients such as amino acids [76] and sugars [77], the utilization of these nutrients by Pl may represent another way to suppress the virulence of Ps. In conclusion, another side effect of Ps rescuing Pl by ITC detoxification might be the suppression of Ps virulence to the plant.

Detoxification of plant specialized metabolites, such as 4MSOB-ITC, may hence be highly relevant in plant-microbe-microbe interactions. We suggest several possibilities of how the Ps virulence factor SaxA may indirectly contribute to plant health by rescuing commensals like Pl. This knowledge contributes to a more holistic understanding of the microbial interactions occurring with significant implications for plant resistance to pathogens and may also contribute to the development of new biocontrol agents.

## Supporting information

Supplementary File DEGs

## Acknowledgements

We thank Dr. Shuaibing Zhang and Prof. Pierre Stallforth for providing us with pEXG2 plasmid. We thank Manuel Velasco Gomariz and Dr. Kathrin Fröhlich for supporting us with their expertise in RNA extraction.

## Data Availability Statement

All data will be made publicly available on figshare before final publication. (https://figshare.com/projects/Pseudomonas_virulence_factor_SaxA_is_a_public_good_that_protects_a_commensal_plant_bacterium_from_a_glucosinolate-derived_host_defence_metabolite/242066)

