## Supplementary Material for "*Pseudomonas* virulence factor SaxA detoxifies plant glucosinolate hydrolysis products, rescuing a commensal that suppresses virulence gene expression"

Supplementary Methods

Supplementary Tables 1-4

Supplementary Figures 1-3

#### Supplementary Methods

Knock-out of *saxA* in *Pseudomonas viridiflava* 3D9

First upstream (UP) and downstream (DW) regions with an overlap of 78 nt and 84 nt with *saxA* were amplified: 5 µL 5x Q5 Reaction Buffer, 0.5 µL 10 mM dNTPs, 1.25 µL 10 µM upA_fwd / downA_fwd, 1.25 µL 10 µM upA_rvs / downA_rvs, 0.25 µL Q5 High-Fidelity DNA Polymerase, 15.75 µL NFW, 1 µL gDNA. Initial denaturation: 98°C for 30 s, 30 cycles: 98°C for 10 s, 65 °C for 5 s, 72°C for 15 s, final extension: 72°C for 2 min, hold at 10°C. Both fragments were cleaned up using the QIAquick PCR & Gel Clean UP Kit (QIAGEN) and eluted in 30 µL 10 mM Tris-HCl pH=8. The UP fragment is 470 nt long and DW fragment is 483 nt long, and this strategy would result in a truncation of 729 nt in the gene *saxA*.

The pEXG2 plasmid was linearized: 20 µL pEXG2 (~1µg), 5 µL 10x rCutSmart Buffer, 1 µL EcoRI-HF (New England Biolabs), 1 µL HindIII-HF (New England Biolabs) and 23 µL NFW were mixed, digested for 60 min at 37°C, and inactivated at 80°C for 20 min. Five 50 µL reactions were combined and 700 µL 100% molecular grade ethanol with 25 µL sodium acetate (3M, pH 6.5) was added to precipitate the linear pEXG2 at -20°C for 4 h. To collect the plasmid, it was centrifuged at 20,000 x g for 40 min, washed with 70% ethanol and resuspended in 30 µL 10 mM Tris-HCl pH=8. To assemble the plasmid the Gibson Assembly Matermix (New England Biolabs) was used and the plasmid and UP and DW regions were mixed 1:2 (molar concentrations). The reaction was incubated at 50°C for 60 min and then stored at 10°C until the transformation.

Chemically competent *E. coli* DH5α cells thawed on ice, 2 µL of the assembled plasmid was added and cells incubated for 30 min on ice. Next, they were heat shocked by exposing them to 42°C for 60 s. After another 5 min on ice, 900 µL LB (10 g/L tryptone, 5 g/L yeast extract, 10 g/L sodium chloride, pH = 7.0±0.2) medium was added followed by an incubation at 37°C, 200 rpm, for 1.5 h. The cells were centrifuged at 3500 x g for 5 min, the supernatant was discarded, and the cells were gently resuspended in 150 µL fresh LB broth which were plated on LB agar supplemented with 15 µg/mL gentamicin (Gent). After 48 h of incubation at 37°C positive colonies were picked and checked for the presence of the correctly assembled plasmid: cell material from individual colonies was resuspended in 0.05 M NaOH and boiled at 95°C for 15 min to extract crude DNA which was used for the subsequent PCR using 8pEX_fwd and 8pEX_rvs primers: 5 µL 5x Q5 reaction buffer, 0.5 µL 10 mM dNTPs, 1.25 µL 10 µM 8pEX_fwd, 1.25 µL 10 µM 8pEX_rvs, 0.25 µL Q5 High Fidelity DNA Polymerase, 15.75 µL NFW, 1 µL template. Initial denaturation: 98°C for 30 s, 30 cycles: 98°C for 10 s, 65 °C for 10 s, 72°C for 25 s, final extension: 72°C for 2 min, hold at 10°C. Using the QIAGEN Miniprep plasmid kit pEXG2-A plasmid was extracted from overnight cultures of *E. coli* DH5α-pEXG2-A in LB+Gent.

Using the same heat shock protocol as mentioned before, chemically competent *E. coli* ST18 cells were transformed with the plasmid and plated on LB plates containing gentamicin and 5-aminolevulinic acid (ALA). After 48 h of incubation at 37°C colonies were screened for the presence of the correct insert using 8pEX_fwd and 8pEX_rvs primers. 2 mL of an overnight culture of *E. coli* ST18-pEXG2-A and 2 mL of *Pseudomonas viridiflava* 3D9 overnight culture were used to inoculate 30 mL of fresh LB+Gent+ALA and LB, respectively. Both were incubated at 200 rpm for 2.5 h at 37°C and 30°C, respectively. Cells were harvested by centrifugation (3,500 x g, 5 min), washed with PBS and finally resuspended in 20 mL PBS. The OD600 was normalized to 0.5 for *E. coli* and 1.0 for *Pseudomonas viridiflava.* For the conjugation the strains were mixed in 3:1, 1:1 and 1:3 ratios (v/v), centrifuged again to recover the cell pellets and resuspended in 100 µL PBS. About 50 µL per mating spot were plated onto LB+ALA plates which were incubated at 30°C. On the next day, bacterial cells were resuspended in 800 µL fresh PBS and 10 µL, 100 µL and 500 µL of the suspensions were plated on LB+Gent plates to select against *E. coli* ST18 and for *Pseudomonas viridiflava* transformants. The transformants were streaked on fresh LB+Gent plates and tested for the absence of *E. coli* contamination (*uidA* primers (*30*)), presence of pEXG2-A plasmid with correct inserts (8pEX primers, Supplementary Table 1). For the counterselection positive colonies were cultured on LB supplemented with 10% sucrose at 30°C overnight.

Finally, the successful homologous recombination and thus deletion of *saxA* was checked using 9gen primers (Supplementary Table 1) which bind to the flanking regions of *saxA* in the genome of 3D9. Mastermix: 1.5 µL 10x Buffer B, 0.6 µL 10 mM dNTPs, 0.3 µL 10 µM 9gen_fwd, 0.3 10 µM 9gen_rvs, 1.5 µL 25 mM MgCl2, 9.65 µL NFW, 0.15 µL Taq Polymerase (Biodeal, Markkleeberg, Germany). Initial denaturation: 95°C for 2 min, 30 cycles: 95°C for 0:30 min, 61°C for 0:30 min, 72°C for 1:30 min, final elongation: 72°C for 5:00 min, cooling to 10°C. The product was sequenced (Eurofins Genomics) and to confirm the loss of function, 4MSOB-ITC and 4MSOB-amine were quantified in the supernatant of an overnight culture.

<u>Details on 4MSOB-ITC and 4MSOB-amine quantification with LC-MS</u>

4MSOB-ITC and its breakdown product 4MSOB-amine in bacterial cultures (n = 3) and non-inoculated medium controls (n = 3) were analyzed on an Agilent 1200 HPLC system (Agilent, Santa Clara, CA, United States) coupled to an API3200 tandem mass spectrometer (AB SCIEX, Darmstadt, Germany). The compounds were separated on an Agilent XDB-C18 column (5 cm × 4.6 mm, 1.8 µm, Agilent, Waldbronn, Germany). The mobile phase consisted of 0.05% (v/v) formic acid in ultrapure water as solvent A and acetonitrile as solvent B, at a flow rate of 1.1 mL/min. The elution gradient was: 0-0.5 min, 3-15% B; 0.5-2.5 min, 15-85% B; 2.5-2.52 min, 85-100% B; 2.25-3.5 min, 100% B; 3.5-3.51 min, 100-3% B, 3.51-6 min, 3% B. The ion spray voltage was maintained at 5500 eV in positive mode. The turbo gas temperature was set to 500 °C, nebulizing gas to 60 psi, drying gas to 60 psi, curtain gas to 35 psi, and collision gas to 3 psi. Details of multiple reaction monitoring (MRM) can be found in Supplementary Tab. 2. Analyst Software 1.6 Build 3773 (AB SCIEX) was used to acquire and process the data.

Details on RNA extraction

The frozen bacterial cell pellets were resuspended in 600 µL lysozyme solution (0.5 mg/ml in TE Buffer, pH=8) and transferred into a 2 mL tube. Samples with the gram-positive Pl were incubated for 5 min at 37°C before adding 60 µL of 10 % SDS. SDS was immediately added to all other samples with only gram-negative Ps. The samples were mixed by inverting the tube and incubated at 64°C for 1-2 min before 66 µL 1 M sodium acetate (pH= 5.2) were added. Next, 750 µL Roti-Aqua-Phenol (Carl Roth, Germany) was added and the samples were incubated at 64°C with constant shaking for 6 min. After 1 min on ice, the samples were centrifuged at 20,000 x g for 25 min at 4°C to separate the phases. The aqueous layer was transferred into phase-lock-tubes (5PRIME Phase Lock Gel heavy; Quantabio, Beverly, Massachusetts, USA) and 750 µL chloroform was added. Then, the samples were shaken for 10 s and incubated for 2-5 min at room temperature. For phase-separation, they were centrifuged at 20,000 x g for 10 min at 4 °C. The aqueous layer was transferred into a new tube and 1.4 mL of a 30:1 mix (ethanol 100 % : 3 M sodium acetate pH=6.5) were added. To recover the RNA/DNA pellet after an overnight incubation at -20°C, the samples were centrifuged at 20,000 x g for 30 min at 4°C. The pellet was washed with 80 % molecular grade ethanol, air-dried and eluted in 30 µL nuclease-free water (NFW) by shaking incubation at 66°C for 5 min. 10-15 µg RNA/DNA were incubated with 5 µL RDD buffer and 1.25 µL DNAse I (QIAGEN) for 10 min at 25°C. The volume was filled up to 100 µL with NFW and 100 µL phenol-chloroform-isoamyl alcohol (25:24:1, Carl Roth) was added into the phase-lock-tube. The samples were shaken for ca. 15 s and incubated at room temperature for 2-5 min. Using another phase-lock-tube the phases were separated after centrifugation at 20,000 x g for 15 min at 12°C. The upper aqueous phase was transferred into a new tube and 300 µL of 30:1 ethanol sodium acetate mix was added to precipitate the remaining RNA overnight at -20°C. On the next day, RNA was recovered by centrifugation at 20,000 x g for 30 min at 4°C, washing with 80 % ethanol, air-drying the pellet and resuspending it at 66°C for 5 min in 30 µL NFW. The concentration of RNA was checked on a Nanodrop spectrophotometer and its presence and quality were checked on an 1.5 % agarose gel. If DNA contamination was observed the samples were treated a second time with DNAse.

### Supplementary Tables

**Supplementary Table 1:** Primer sequences used in this study.

| Primer name | Primer sequence | Binding site and product length(s) | Reference |
|---|---|---|---|
| upA_fwd | acgagccggaagcataaatgtaaagcaCCGAACGCCTCCAG TTGATG | Amplification of upstream (UP) fragment | This study |
| upA_rvs | taccagccctgccTTACGTTCCTACGTCAAGCAG | | |
| downA_fwd | acgtaggaacgtaaGGCAGGGCTGGTATCAGCGAAA G | Amplification of downstream (DW) fragment | This study |
| downA_rvs | cacccgtggaaattaattaaggtaccgAATGCAGAGGCAGG CCCGAG | | |
| 8pEX_fwd | ACGGCAGGTAAGCTAATTCCAC | Binds to flanking region of inserts in pEXG2 plasmid | This study |
| 8pEX_rvs | CCTCAACGACAGGAGCACGA | | |
| 9gen_fwd | TGCACATATGGCTCATCGCA | Binds to flanking regions of *saxA* gene in genome of 3D9 | This study |
| 9gen_rvs | AATGCGTCGTCGCTTCCT | | |
| FWD_uidA | AACAGGTGGTTGCAACTGGA | Binds to β-d-glucuronidase gene in *E. coli* | (*31*) |
| REV_uidA | TTGCTGAGTTTCCCCGTTGA | | (*31*) |

**Supplementary Table 2. Details of the analysis of 4MSOB-amine and 4MSOB-ITC by LC-MS/MS.** Compounds were measured using an Agilent HPLC 1200/API3200 (AB SCIEX) instrument in positive ionisation mode. Abbreviations are: Q1, selected *m/z* of the first quadrupole; Q3, selected *m/z* of the third quadrupole; RT, retention time; DP, declustering potential (V); and CE, collision energy (V).

| **Q1** | **Q3** | **RT (min)** | **compound** | **DP** | **CE** |
|---|---|---|---|---|---|
| 136 | 72 | 0.5 | 4MSOB-amine | 26 | 17 |
| 178 | 114 | 2.6 | 4MSOB-ITC | 60 | 13 |

**Supplementary Table 3: *saxA* and *saxB* expression in Ps.** Log2FoldChanges (L2FC) and adjusted p-values (Wald test with Benjamin-Hochberg correction) of DESeq2 analyses of *saxA*, *saxB*, *saxC* genes.

| Gene name / Locus Tag | **Condition** | **Effect** | **L2FC** | **padj** |
|---|---|---|---|---|
| ***saxA* / MIFLLO_11060** | Psco | ITC | 0.523789 | 0.00116082 |
| | Psmono | ITC | 0.629637 | 0.00569671 |
| | PsnoITC | Partner | 0.171169 | 0.35224 |
| | PsITC | Partner | 0.0511512 | 0.736944 |
| ***saxB* / MIFLLO_11060** | Psco | ITC | 0.368971 | 0.0216534 |
| | Psmono | ITC | 0.441532 | 0.0451459 |
| | PsnoITC | Partner | -0.0891716 | 0.600876 |
| | PsITC | Partner | -0.196045 | 0.13862 |
| ***saxC* / MIFLLO_11070** | Psco | ITC | -0.0450266 | 0.641786 |
| | Psmono | ITC | 0.0502467 | 0.705504 |
| | PsnoITC | Partner | 0.805898 | 2.14999e-8 |
| | PsITC | Partner | 0.453799 | 0.000257485 |

**Supplementary Table 4: *saxF* expression in Ps.** Log2FoldChanges (L2FC) and adjusted p-values (Wald test with Benjamin-Hochberg correction) of DESeq2 analyses of *saxF*-associated efflux pump genes (Multidrug efflux pump subunit AcrB, with COG0841 annotation).

| Locus Tag | Condition | Effect | L2FC | padj |
|---|---|---|---|---|
| MIFLLO_05915 | Psco | ITC | 0.0214242 | 0.915358 |
| | Psmono | ITC | 0.118351 | 0.370856 |
| | PsnoITC | Partner | 0.584841 | 3.50777e-8 |
| | PsITC | Partner | 0.468382 | 1.93891e-11 |
| MIFLLO_11330 | Psco | ITC | 0.000448884 | 0.998677 |
| | Psmono | ITC | 0.0290503 | 0.832323 |
| | PsnoITC | Partner | 0.944021 | 4.28782e-10 |
| | PsITC | Partner | 0.832762 | 6.43417e-12 |
| MIFLLO_13095 | Psco | ITC | -0.00464959 | 0.984911 |
| | Psmono | ITC | 0.0523958 | 0.697142 |
| | PsnoITC | Partner | -0.321806 | 0.00606728 |
| | PsITC | Partner | -0.464192 | 1.33456e-7 |
| MIFLLO_14945 | Psco | ITC | -0.000697519 | 0.997347 |
| | Psmono | ITC | 0.0491537 | 0.641425 |
| | PsnoITC | Partner | 0.133161 | 0.431736 |
| | PsITC | Partner | -0.0915416 | 0.577871 |
| MIFLLO_15950 | Psco | ITC | -0.0216916 | 0.885046 |
| | Psmono | ITC | 0.0252796 | 0.848564 |
| | PsnoITC | Partner | 0.133161 | 0.431736 |
| | PsITC | Partner | 0.120148 | 0.397794 |
| MIFLLO_18075 | Psco | ITC | 0.0126581 | 0.928562 |
| | Psmono | ITC | 0.0310502 | 0.809603 |
| | PsnoITC | Partner | -0.00689545 | 0.970914 |
| | PsITC | Partner | -0.0620324 | 0.692812 |
| MIFLLO_21925 | Psco | ITC | -0.0351024 | 0.785644 |
| | Psmono | ITC | -0.00701026 | 0.977402 |
| | PsnoITC | Partner | 0.0381757 | 0.790674 |
| | PsITC | Partner | -0.0458315 | 0.694308 |
| MIFLLO_25910 | Psco | ITC | -0.0158039 | 0.927966 |
| | Psmono | ITC | -0.0865286 | 0.46315 |
| | PsnoITC | Partner | -1.47936 | 1.70152e-16 |
| | PsITC | Partner | -1.26254 | 7.3265e-47 |

**Supplementary Table 5. Reaction of Pl to 4MSOB-ITC.** $Log_2$FoldChanges (L2FC) and adjusted p-values (Wald test with Benjamin-Hochberg correction) of DESeq2 analyses of Pl's response to 4MSOB-ITC in monoculture.

| Locus Tag | L2FC | padj | protein annotation from whole genome |
|---|---|---|---|
| ACEBMG_12325 | -2,725 | 6,71E-08 | Aldehyde dehydrogenase B |
| ACEBMG_11515 | -2,715 | 2,95E-16 | hydroxymethylglutaryl-CoA reductase, degradative |
| ACEBMG_12320 | -2,671 | 1,24E-51 | UDP-glucose 4-epimerase |
| ACEBMG_11450 | -2,288 | 5,30E-56 | PepSY domain-containing protein |
| ACEBMG_09335 | -2,271 | 1,74E-37 | 3-oxopropanoate dehydrogenase |
| ACEBMG_17620 | 2,003 | 1,35E-57 | Alcohol dehydrogenase |
| ACEBMG_12995 | 2,053 | 8,79E-77 | NADPH dehydrogenase |
| ACEBMG_15675 | 2,128 | 7,56E-71 | N-ethylmaleimide reductase |
| ACEBMG_17225 | 2,129 | 4,12E-10 | HxlR family transcriptional regulator |
| ACEBMG_16910 | 2,148 | 3,29E-58 | HTH-type transcriptional regulator CmtR |
| ACEBMG_17655 | 2,191 | 1,51E-20 | Alpha/beta fold hydrolase |
| ACEBMG_15290 | 2,194 | 1,96E-83 | NADPH:quinone reductase |
| ACEBMG_03985 | 2,224 | 1,56E-89 | Nitroreductase |
| ACEBMG_17650 | 2,255 | 1,40E-57 | Pyrroline-5-carboxylate reductase catalytic N-terminal domain-containing protein |
| ACEBMG_17080 | 2,280 | 2,06E-23 | Copper chaperone |
| ACEBMG_03025 | 2,666 | 1,29E-27 | AraC family transcriptional regulator |
| ACEBMG_01540 | 2,672 | 4,88E-37 | NAD-dependent epimerase/dehydratase family protein |

### Supplementary Figures

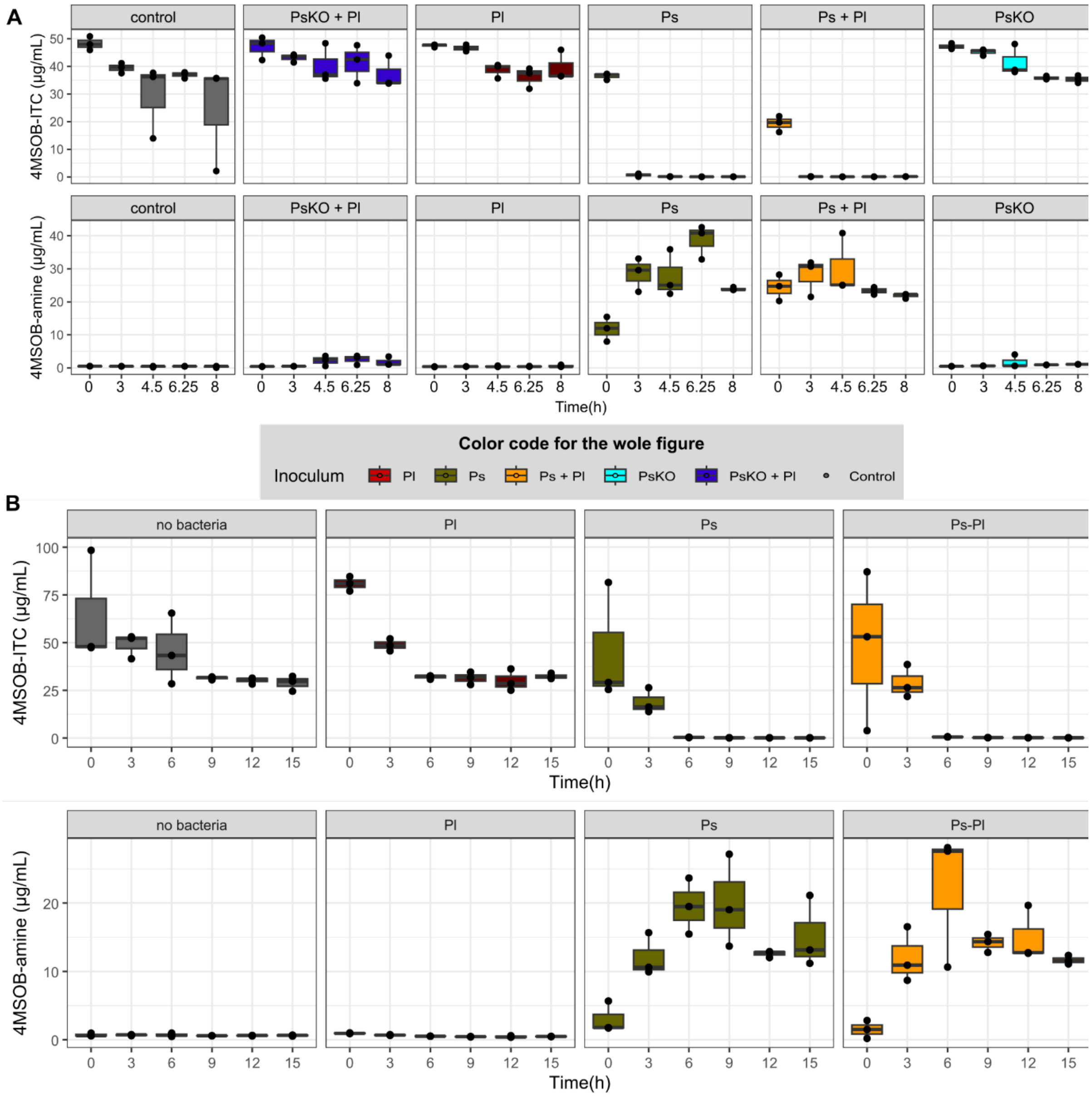


**Supplementary Figure 1: 4MSOB-ITC and 4MSOB-amine quantification with HPLC-MS over time.** 4MSOB-ITC and -amine were measured in bacterial supernatants over the course of 8 h (**A**) and 15 h (**B**). Starting concentrations were either 60 µg/mL (A) and 30 µg/mL (B) 4MSOB-ITC. Controls were not inoculated with bacteria but with sterile medium, all samples and controls n=3.

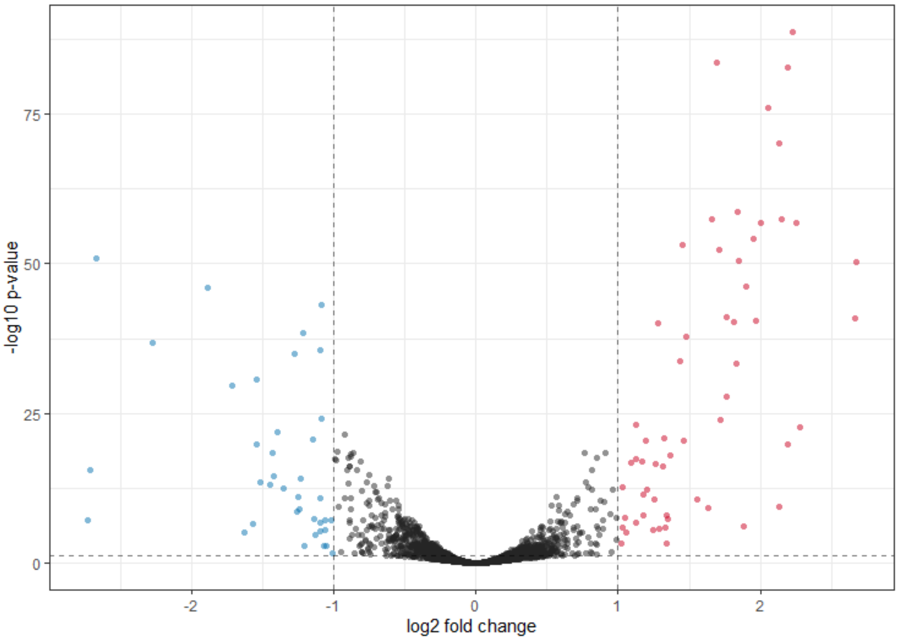


**Supplementary Figure 2: Pl's reaction to 4MSOB-ITC in monoculture.** Volcano plot of significant DEGs (adjusted p-value < 0.05, |L2FC| >1). Significant genes are shown in blue (downregulated) or red (upregulated).

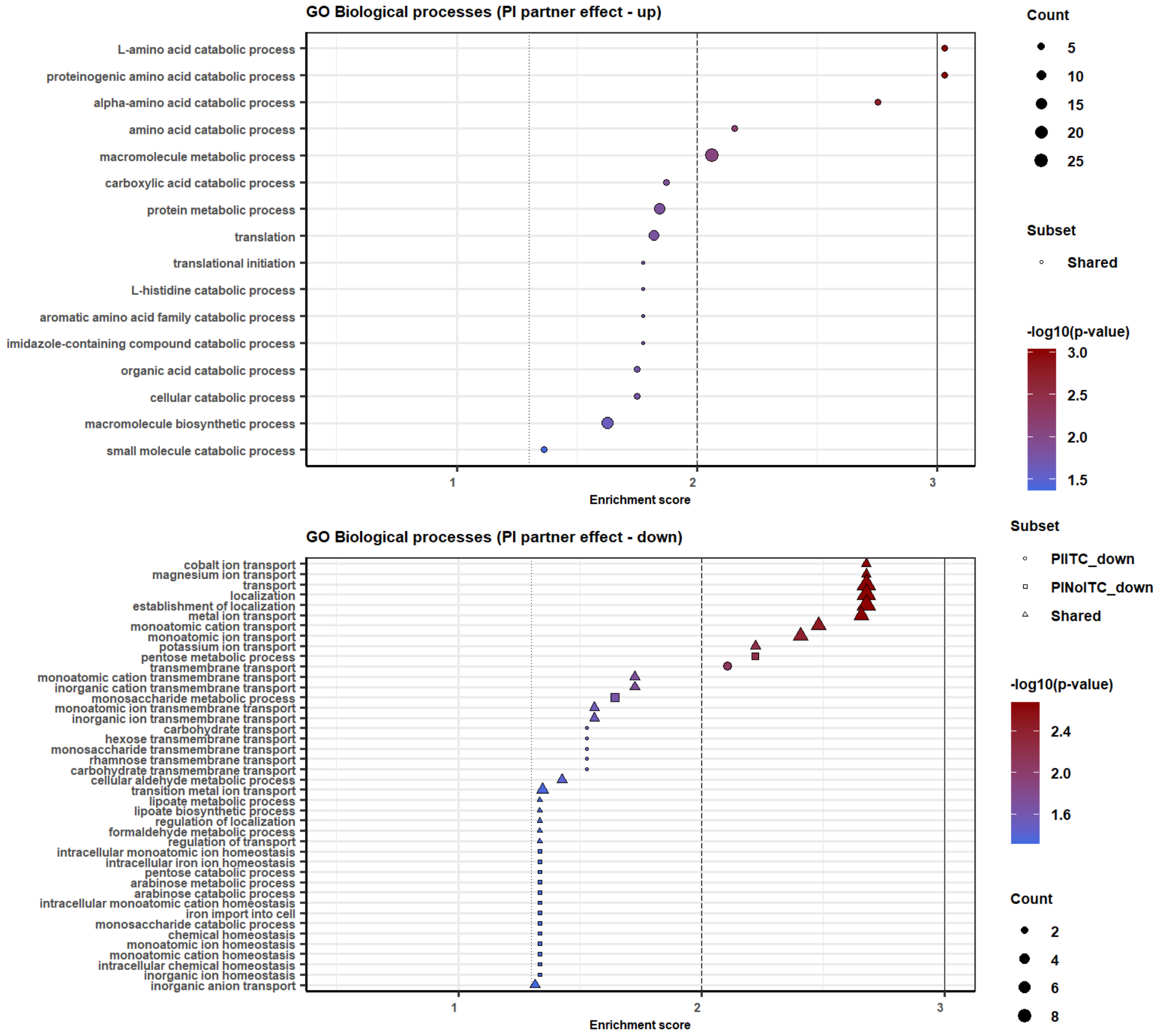


**Supplementary Figure 3: Ps's reaction to Pl dependent on 4MSOB-ITC exposure.** (A) Enriched GO terms for up-regulated Ps DEGs. (B) Enriched GO terms for down-regulated DEGs. The shape shows whether a term was enriched only with or without 4MSOB-ITC or independent of it. The color indicates the p-value of the Fisher test, horizontal lines depict p=0.001, p=0.01 and p=0.05. The size shows how many GO terms were identified for each biological process.
